# One-to-one Benefit provided by Antioxidants to cultured skin Fibroblasts from Friedreich Ataxia patients

**DOI:** 10.1101/2024.04.25.591088

**Authors:** Paule Bénit, Malgorzata Rak, Pierre Rustin

## Abstract

With the hope of better understanding and characterizing the dramatic differences in antioxidant response of human cells carrying mutations in the frataxin gene responsible for Friedreich ataxia (FRDA), we studied primary cultures of skin fibroblasts derived from five FRDA patients harboring different large expansion sizes in the frataxin gene. Because oxidative stress was previously established to be instrumental in FRDA, among the many enzymes likely to modulate sensitivity to oxidative stress, we selected some previously shown to play a critical role in this stress. However, we failed to identify a consistent clue to order individual cultures based on these measurements. We therefore focused our study on cell death of FRDA fibroblasts and its modulation by different antioxidants under culture conditions exacerbating sensitivity to oxidative stress. Under conditions that force the cells to rely on mitochondrial activity, we observed significant yet variable FRDA cell proliferation. These conditions were thereafter used to screen the effectiveness of a set of antioxidant molecules targeting different steps of the pro-oxidant cascade previously documented in FRDA, *i.e.* Uridine, Pyruvate, and Pioglitazone, to prevent or slow down cell mortality. We observed a surprising variability of response to antioxidant molecules even under the similar, controlled conditions used to culture patient’s fibroblasts. We conclude that the specific response of each individual already discernable at cellular level may well play an important role in the frequent difficulties encounter to reach firm conclusions when testing the capacity of antioxidants to counteract the consequences of frataxin depletion, including in clinical trials.

## Introduction

Friedreich ataxia (FRDA) is an autosomal recessive hereditary mitochondrial disease (1/30,000 births) ranked among neurological diseases [1,2]. Often detected during adolescence or in young adults, the onset may also occur in early childhood and even in toddlers [3], or conversely the disease may arise much latter, up to 50 years of age or over [4–6]. Some patients endure a rapid and severe progression while others experience a very slow, somewhat erratic course of their disease. Patients with early-onset disease (before 15 years old) generally worsen faster than others. Documented neurological features of FRDA are progressive trunk and limbs mixed cerebellar and sensory ataxia, associated with dysarthria, pyramidal signs. Beside neurological symptoms, skeletal deformities (scoliosis, foot deformities) and cardiomyopathy are found in most patients, who also have an increased frequency of diabetes. Visual, hearing impairment, dysphagia and bladder dysfunction are also encountered. Neurological signs may be subtle at onset with hypertrophic cardiomyopathy being the prominent symptom initially detected. Conversely, the heart can remain asymptomatic throughout life in other patients.

Most of the cases (>95%) result from abnormal GAA-triplet expansions of different length on both alleles of the first intron of the Frataxin gene located on chromosome 9 [7]. These expansions reduce gene transcription resulting in low Frataxin levels (30 to 2% of control), an essential mitochondrial protein involved iron-sulfur cluster (ISC) biogenesis [8]. In the heart micro biopsies of a series of FRDA patients, a generalized deficiency of mitochondrial respiratory chain (RC) ISC-containing proteins (ISP) was evidenced together with defects of both cytosolic and mitochondrial ISC-containing aconitases [9]. Similarly, in a number of organisms studied in laboratories, Frataxin deficiency results in impaired synthesis of ISC in the mitochondria [8]. In cultured human fibroblasts grown under standard conditions, Frataxin depletion does not significantly affect the activity of ISC-containing enzymes, either involved in the RC or the Krebs cycle [9]. Yet, it often causes a slight overproduction of superoxides revealed by an elevation of the superoxide-inducible SOD activity [10] and a low content in reduced glutathione [11–13]. These cells simultaneously display a frequent hypersensitivity to external oxidizing agents [14]. This increased susceptibility to a number of oxidative agents, including oxygen itself [15], is indeed observed in a number of cell types, animals, including humans, where Frataxin gene is mutated and where poor handling of oxidizing pressure is presumably instrumental in disease progression [15–17]. The prominent role played by oxidant stress can be exacerbated when a long expansion of the Frataxin gene additionally disturbs the transduction of the neighboring gene, PIP_5_K_1_β, coding the PIP_5_K protein [18,19]. Impairing the transcription on the PIP_5_K-encoding gene destabilizes cell actin-network and causes major disorganization of actin-bound Keap_1_/Nrf_2_ antioxidant signaling proteins ultimately resulting in impaired antioxidant cell defenses [10].

Thus, whether considering the genetic aspects (*e.g.,* size of the expansion in the Frataxin gene or identity of the genes involved, *i.e.,* Frataxin, PIP_5_K_1_β), the biochemical aspects (*e.g.,* damage to soluble ISP, to RC components or to antioxidant systems), the clinical signs and the sequence of their apparition, a very contrasting picture between patients emerges. Indeed, this inter-individual variability hallmarks the disease in the foreground [3,20–31].

Considering the context of this rare, slowly progressive disease, possibly resulting from the combination of several mechanisms, with rather unpredictable steps, the almost insurmountable difficulty to find a drug or a procedure equally effective for all patients becomes understandable. For the same reasons, organizing, interpreting therapeutic trials, making predictions about chances of therapeutic success, all have turned out to be true nightmares. Thus, not surprisingly, 26 years after the discovery of the genetic bases of the disease in 1996 [7], therapeutic trials have proved to be essentially inconclusive. So far statistical power of trials remains insufficient to reach firm conclusion, despite the frequent occurrence of sub-groups of patients positively responding to the treatment when none of the individuals receiving a placebo did [32]. A much better identification and understanding of the mechanisms that could explain the great variability observed between individuals appear therefore essential.

In the present study, we focused on the oxidative stress and its management targeting few steps of the Nrf2 pathway in cultured skin fibroblasts. In 2009, we identified this pathway as defective in FRDA in association with abnormalities of fibroblasts actin network [10]. The defect of Nrf2 antioxidant pathway was found more severe in case of impaired expression of the Frataxin-adjacent PIP5k-encoding gene caused by particularly large expansion in the Frataxin-gene [19]. This pioneer work paved the way to identify molecules targeting this pathway, (Pioglitazone, Leriglitazone, Omaveloxolone). Noticeably this latter drug, in the absence of any available therapy, is the only one so far approved by the FDA for FRDA (https://www.fda.gov/drugs/news-events-human-drugs/fda-approves-first-treatment-friedreichs-ataxia), since we and other have documented its frequent occurrence and partly delineated the putative underlying mechanism. Using precisely the five cell lines derived from FRDA patients for which we previously gather a set of molecular and biochemical data, we here underlines the one-to-one response to three substances known to exert antioxidant effects by distinct mechanisms (Fig. 1) : Uridine, acting particularly on the antioxidant signaling pathway of SOD1 (cytosolic and mitochondrial intermembrane location) [33] through the glyoxalase DJ-1 [34,35]; Pyruvate, a potent hydrogen peroxide detoxifying agent through a non-enzymatic reaction specific to dicarboxylic acids [36]; Pioglitazone acting on antioxidant defenses signaling, in particular of mitochondrial SOD2, via the Nrf2 transcription factor to rescue cells mortality in FRDA [37].

**Figure 1.**
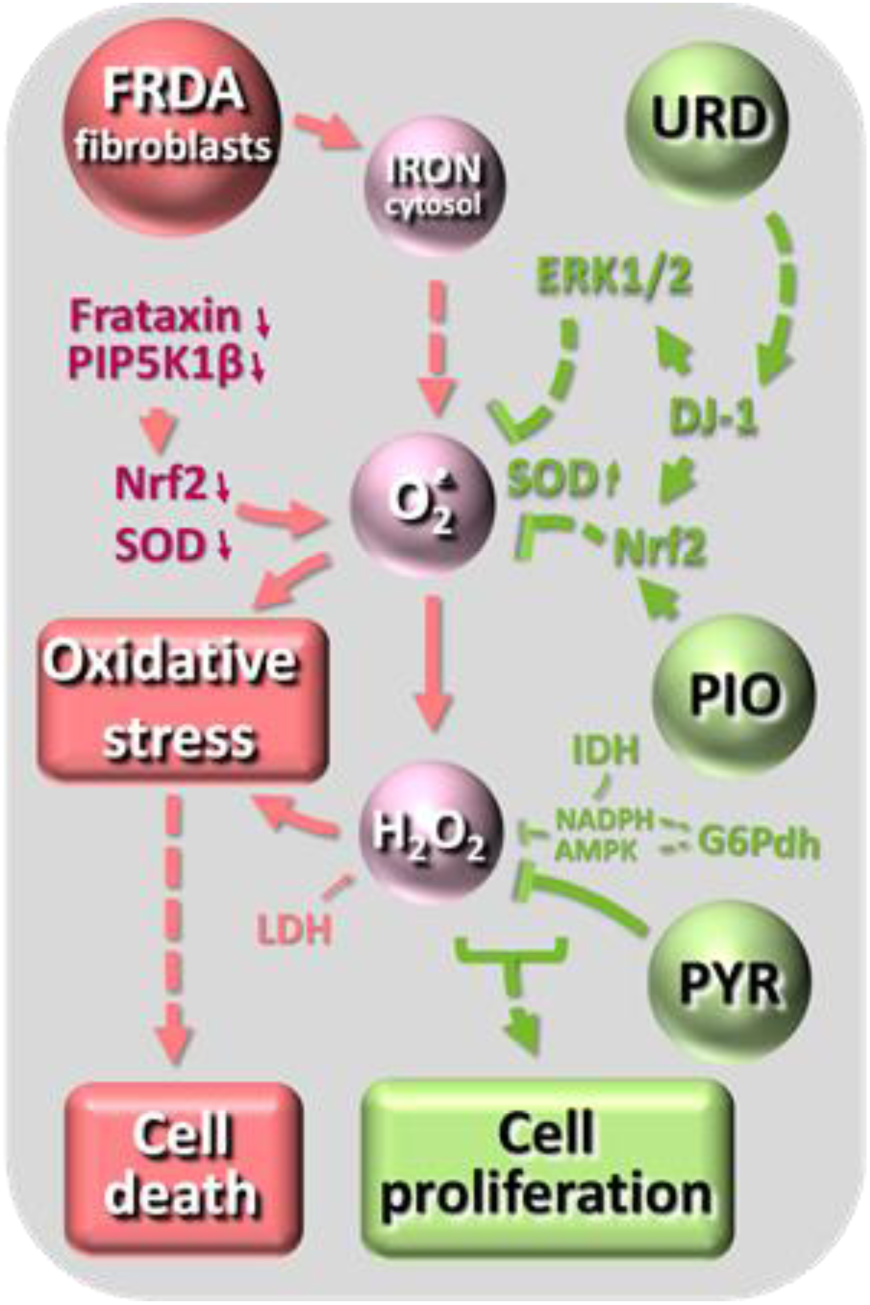
A very simplified diagram showing the action sites of the antioxidant molecules tested for their effect on fibroblast growth. On the left, the potential consequences of Frataxin gene expansion, in the middle the oxidative stress cascade, on the right, in green, the action sites of Uridine (URD), Pioglitazone (PIO) and Pyruvate (PYR) on the oxidative stress cascade. PIP_5_K_1_β, phosphatidylinositol-4-phosphate 5-kinase type 1 β; ERK1/2, extracellular signal-regulated kinase 1/2 cascade; DJ-1, a glyoxalase [34] [35]) also known as Parkinson protein 7 (Park7); Nrf_2_, nuclear factor (erythroid-derived 2)-like 2; SOD, superoxide dismutase.

## Materials and Methods

### Human skin fibroblasts

Cultured fibroblasts were derived from skin biopsies obtained from five FRDA patients (FRDA1 to 5), and four anonymous healthy individuals, all from European ancestry. Written informed consent for research was obtained from patients and/or family members according to protocols in accordance with Robert Debré Hospital ethical committees (FRDA patient; 2011; Paris, France). The FRDA fibroblasts were from patients harboring a biallelic long GAA expansions (>2.6 kb) in the Frataxin (FXN) gene. These expansions were previously shown to impair the transcription of the adjacent PIP_5_K_1_β gene [18]. Compared to controls residual PIP_5_K_1_β mRNA was 58, 50, 43, 33 and 24% respectively [18] for patients noticed 1 to 5 in our study.

Cells were grown in T75 flasks in 10 ml Dulbecco’s modified Eagle’s minimal essential medium (DMEM) containing 4.5 g/l Glucose, 4 mM Glutamine (Glutamax; Gibco Thermo Fisher Scientific), 2 mM Pyruvate and 200 μM Uridine (permissive complete medium, thereafter referred as CM) or lacking Glucose, Pyruvate and Uridine but containing 4 mM Glutamine (non-permissive medium, thereafter referred as SC for stressing condition) or SC *plus* Uridine (SCU), or SC *plus* Pyruvate (SCP) or both Uridine and Pyruvate (SCUP). All media were supplemented with 10% fetal calf serum, 100 U/ml penicillin and streptomycin each. Cell pellets (1,500 *g* for 5 min) were kept frozen (−80°C) for ulterior biochemical enzyme assays studies.

Cells from a confluent culture (75-cm^2^ flask) were trypsinized and seeded in 4×75-cm^2^ flasks (20–25% confluency) containing permissive CM medium. After allowing the culture to stand for one night, the medium was changed to the desired conditions, *i.e.,* CM medium, SC medium, or SCU, SCP, SCUP, SC or SCU *plus* DMSO (0.1 %) or 10 µM Pioglitazone in DMSO (Pioglitazone hydrochloride, Sigma, France). The number of cells was estimated from random image sampling using the ImageJ processing program for 14 days without changing the medium. Noticeably, using brand new DMSO solutions is imperative as upon opening and several months of storage the solution becomes progressively pro-oxidative and deleterious for cultured cells. Due to its broad spectrum of solubility, DMSO still remains an often-irreplaceable solvent, *e.g.,* for the development process of new antioxidants with neuroprotective properties [38,39].

### Microscopy

Representative random phase-contrast photographs were taken on an LSM 5 Ex[1]citer optic microscope (Eclipse TE300 Nikon, France) (×4). The area of cells cultured in CM medium was estimated from random image sampling using the ImageJ processing program.

### Enzyme measurements

Superoxide dismutase (total SOD) (EC 1.15.1.1) and catalase (EC 1.11.1.6) activities of cells cultured in CM and SC medium were measured either spectrophotometrically (SOD) or from oxygen release (catalase) at 37°C on frozen cell pellets according to previously described protocols [40].

The activity of various enzyme implicated in glucose metabolism and NADPH production: NADP^+^ Glucose-6-Phosphate dehydrogenase (G6PDH) (EC 1.1.1.49), NAD^+^ lactate dehydrogenase (LDH) (EC 1.1.1.27), NADP^+^-dependent isocitrate dehydrogenase (IDH1-2) (EC 1.1.1.42), NAD^+^ glutamate dehydrogenase (GDH) (EC 1.4.1.2) activities were measured in CM conditions. All activities were spectrophotometrically measured (Cary 60, Varian, Agilent Technologies) on frozen-thawed cell pellets, at 37°C according to published protocols [41]. Protein was estimated by the Bradford assay.

### Statistics

Data are presented as mean ± SD for all experiments. Statistical significance was calculated by standard unpaired *t*-test or one-way ANOVA with Bonferroni post-test correction for more than two conditions. A *p* ≤ 0.05 was considered statistically significant (GraphPad Prism 5) and noticed with one asterisk, *p* values less than 0.01 with two asterisks, *p* values less than 0.001 with three asterisks.

## Results

### Stressing conditions for studying the effect of antioxidants

We first observed that, as previously reported [10], most FRDA fibroblasts cultured in medium containing high Glucose and Glutamine (providing substrates for both cell proliferation and antioxidant defenses), *plus* Uridine and Pyruvate increase their basal level of antioxidant defenses. This is reflected by a slight but significant increased level of basal SOD activity in the FRDA patient’s fibroblasts. This difference is abolished in stressing condition medium (SC) containing only glutamine as carbon source (Table 1). No similar increase was observed for catalase activity whatever the culture condition used. Noticeably the fibroblasts from one of the FRDA patient displayed a particularly low catalase activity. Similarly, even when cells were cultured in CM condition, G6PDH, NADP^+^ IDH, NAD^+^ GDH and LDH activities were found unchanged (Table1).

**Table 1.** Group *versus* individual enzyme activities measured in control or skin fibroblasts cultures from five FRDA patients. As indicated, superoxide dismutase and catalase have been measured after four days of cell culture either in complete medium or under stressing conditions (media described in details under material and methods). * significantly different from controls.

| Enzyme | Control<br>fibroblasts<br>(n=4; 3<br>assays each) | FRDA<br>fibroblasts<br>(n=5; 3<br>assays each) | FRDA1 | FRDA2 | FRDA3 | FRDA4 | FRDA5 |
| --- | --- | --- | --- | --- | --- | --- | --- |
| **Total SOD**<br>(U/mg/prot) |  |  |  |  |  |  |  |
| - Complete medium | 3.7 ± 0.5 | 4.9 ± 0.5* | 4.4 ± 1.8<br>(n=3) | 4.4 ± 1.4<br>(n=4) | 5.3 ± 1.5<br>(n=3) | 5.2 ± 2.7<br>(n=3) | 5.2 ± 1.7<br>(n=3) |
| - Stressing conditions | 3.7 ± 1.0 | 3.8 ± 0.4 | 4.2 ± 0.4<br>(n=4) | 4.0 ± 0.5<br>(n=3) | 4.1 ± 0.8<br>(n=3) | 3.6 ± 1.0<br>(n=3) | 3.2 ± 0.3<br>(n=4) |
| **Catalase**<br>(nmol /mg/prot) |  |  |  |  |  |  |  |
| - Complete medium | 377±118 | 434 ±215 | 550±137<br>(n=3) | 534±136<br>(n=3) | 411±150<br>(n=3) | 71±24*<br>(n=3) | 603±112<br>(n=4) |
| - Stressing conditions | 485±90 | 584±73 | 627±105<br>(n=3) | 623±152<br>(n=4) | 649±113<br>(n=3) | 172±67*<br>(n=3) | 550±168<br>(n=3) |
| Glucose-6-phosphate dehydrogenase<br>(nmol/min/mg prot) | 117±15 | 96±20 | 125±30<br>(n=9) | 78±11<br>(n=8) | 99±23<br>(n=7) | 74±23<br>(n=7) | 100±30<br>(n=7) |
| NADP<sup>+</sup> Isocitrate dehydrogenase<br>(nmol/min/mg prot) | 16.3±2.2 | 15.7±3.5 | 18.9±6.6<br>(n=7) | 13.6±4.3<br>(n=7) | 12.8±5.6<br>(n=7) | 13.3±5.2<br>(n=7) | 22.3±3.0<br>(n=7) |
| NAD<sup>+</sup> Glutamate dehydrogenase<br>(nmol/min/mg prot) | 9.8± 4.1 | 8.9±3.1 | 7.7 ± 0.2<br>(n=3) | 11 ± 0.1<br>(n=3) | 9.8±1.0<br>(n=3) | 5±0.8<br>(n=3) | 8.7±2.9<br>(n=3) |
| Lactate dehydrogenase<br>(nmol/min/mg prot) | 2306±750 | 3103±527 | 2470±264<br>(n=3) | 3793±504<br>(n=3) | 2600±656<br>(n=3) | 3234±168<br>(n=3) | 2726±605<br>(n=3) |

Finally, we noticed that the area of FRDA patient’s cells was significantly lower than that of controls cells which can be attributed to previously described cytoskeletal structure disorganization possibly resulting from inconspicuous oxidative stress (Supplemental Fig. 1).

To select stressing culture conditions (SC), we first evaluate cell proliferation in rich (CM) *versus* stringent (SC) culture medium (2 FRDA patients and 2 controls). Of note, FRDA fibroblasts actively grow under standard conditions, echoing the absence of significant enzyme defect in these cells [9]. Thereafter referred as the complete culture medium, CM consisted of a Glucose-rich DMEM medium added with Glutamine, Pyruvate and Uridine (CM). In contrast, SC medium was limited to Glutamine as a carbon source, and devoid of any molecule possibly providing an indirect antioxidant reservoir. After 14 days of culture, cell counting revealed a significant difference between control and FRDA fibroblasts: while the growth of control cells in SC medium (Fig. 2) was not significantly affected, the FRDA fibroblasts hardly survived 14 days of culture (at most 25% compared to CM condition).

**Figure 2.**
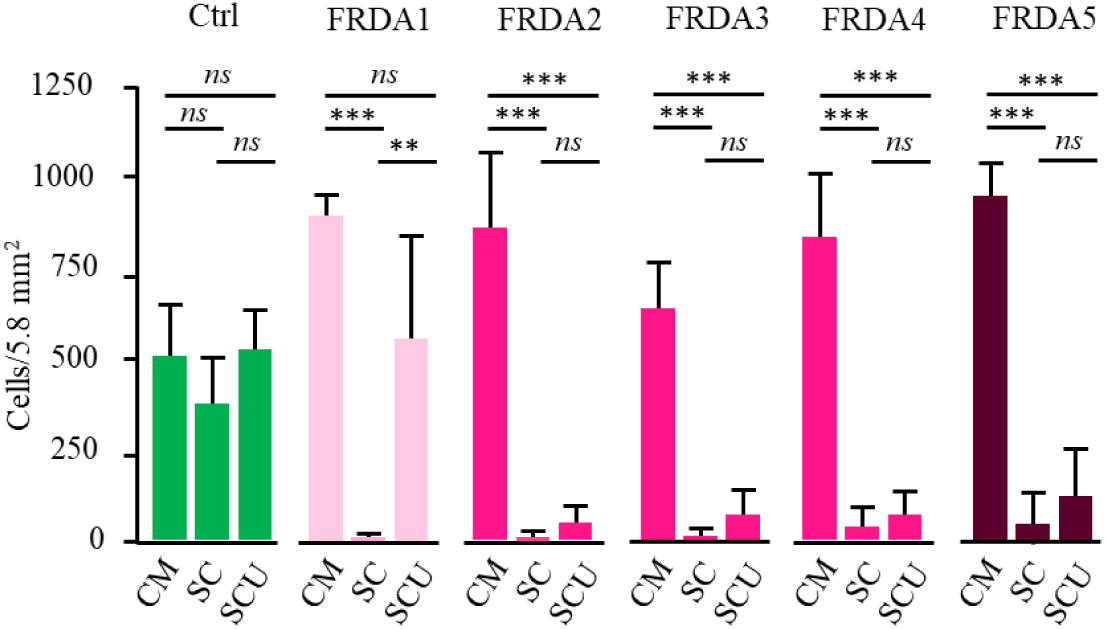
Proliferation of control (n=4; green) and FRDA fibroblast cells (FRDA1 to 5; red) in complete medium (CM), under stressful conditions (SC) and under SC in the presence of Uridine (SCU). Red intensity of the graphs for FRDA cells corresponds to the PIP_5_K_1_β RNA decrease inversely proportional to the extent of the expansions in the Frataxin gene (see Supplemental Table). *ns,* non-significant

Noticeably, a limited, yet significant, cell mortality (50%) could be observed in one control under these quite harsh culture conditions. Addition of Uridine fully preserved cell growth in all control cells. We therefore used SCU medium for studying the effect of Pioglitazone in next experiments (Fig.4). We also verified that none of these cells endured impaired use of Pyruvate. Such impairment would result in Lactate accumulation in case of a RC dysfunction as it has been reported in several human tissues with Frataxin mutation. After 14 days of culture the color of the medium remained unchanged indicating that no significant medium acidification and Lactic acid accumulation (not shown).

### Antioxidant effect of Uridine

SOD being a key limiting enzyme controlling oxidative stress in FRDA, we first selected Uridine as an antioxidant shown to positively act on SOD enzyme expression. Both the manganese SOD_2_- and the copper-zinc SOD_1_-induction by Uridine occurs through the Nrf2-signaling pathway [42,43]. Moreover, SOD_1_-induction by Uridine also occurs through the Elk/Erk pathway. Noticeably the SOD_1_ isoform is present in both the cytosol and the mitochondrial inter-membrane while SOD_2_ is confined to the mitochondrial matrix space. In SCU medium, control fibroblasts readily and similarly proliferate. In contrast, SCU medium did not provide protection against cell death in 4 of the 5 FRDA fibroblasts (Fig. 2). Nevertheless, this medium still allowed a significant growth for cells from patient 1 (FRDA1) (35 % of cell death). FRDA1 patient is distinguished from the other four FRDA patients by harboring the smallest expansion in the Frataxin gene with a limited effect on the transcription of neighboring genes (Table. 2). We previously showed that very large expansion cause silencing of both the FXN and PIP_5_K_1_β genes [18]. Loss of PIP_5_K function results in destabilization of the actin network and impaired Keap_1_-Nrf_2_ signaling of superoxide dismutase (SOD). This resulted in cells becoming highly sensitive to superoxides [10]. Accordingly, these cells (FRDA2) were previously shown to be rapidly killed by exposure to even low doses of distal Q-binding site inhibitors of complex II (SDHI fungicide) due to slight overproduction of superoxides [44].

### Antioxidant effect of Pyruvate

We next tested the effect of Pyruvate known, as other keto acids, to promptly react with hydrogen peroxides [45]. Compared to Uridine (SCU; Fig. 2), Pyruvate alone provides a significantly better protective effect for FRDA fibroblasts (SCP; Fig. 3). However, a significant mortality was still observed, especially substantial for FRDA5. Patient 5 distinguished from the other four FRDA patients by the largest expansion in the Frataxin gene (Table. 2), previously shown to impair PIP_5_K_1_β transcription. This indicated that impaired SOD signaling in this patient’s cells could not be fully overcome by peroxide trapping by Pyruvate.

**Figure 3.**
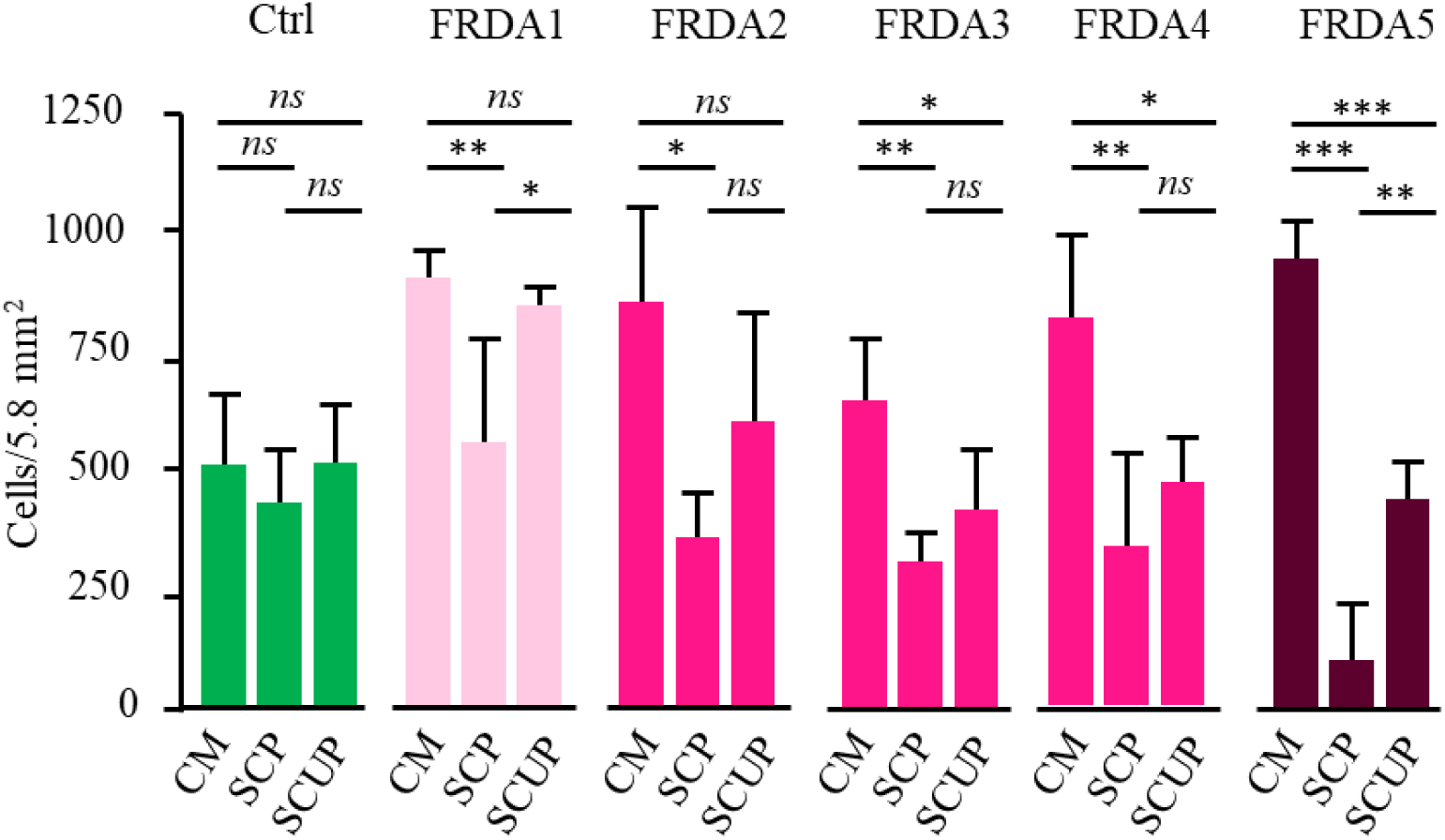
Proliferation of control (n=4; green) and FRDA fibroblast cells (FRDA1 to 5; red) in complete medium (CM), under stressful conditions in the presence of Pyruvate (SCP) or in the presence of Uridine plus Pyruvate. Color code as described in Fig. 2 legend. *ns,* non-significant

### Protecting effect provided by the simultaneous presence of both Uridine and Pyruvate

The combination of the two antioxidants, Uridine and Pyruvate provides a quite significant protection against cell death for four of the five FRDA fibroblasts (SCUP; Fig. 3). Noticeably, the fibroblasts of patient 5, whose growth remained severely affected despite the presence of Pyruvate alone, benefit from the simultaneous presence of Pyruvate and Uridine.

### Probing the antioxidant effect of Pioglitazone

The peroxisome proliferator-activated receptor γ (PPARγ) agonist Pioglitazone was selected because it has been shown to possibly enhance the transcription of Nrf_2_, and several target genes involved in cellular antioxidant response. Noticeably, Pioglitazone has been reported to reduce disease progression in a subset of the neurologically affected mice, the so-called *Harlequin* mice. As patients with FRDA disease, this mouse possesses a heterogeneous genetic background since resulting from a non-intended proviral insertion in the X-linked Aifm1 locus [46]. This causes a 90 to 35% reduction of Aifm1 protein according to tissues [47]. This leads to a time- and tissue-dependent respiratory chain complex I deficiency which classified this affection as a mitochondrial disease similarly to FRDA. Similarity with FRDA, *Harlequin* mouse phenotype is also marked by the spectacularly inconsistent nature and severity of symptoms, including muscle, neurological, ocular, and cardiac symptoms [48]. Individual responders to Pioglitazone were noticed in the *Harlequin* mice [49].

As shown in Figure 4, Pioglitazone afforded no protection against cell death when cells were grown under stressing conditions even when Uridine was added to lighten the stress pressure. This lack of effect indicates that in the fibroblasts from this set of patients the antioxidant defenses pathway targeted by Pioglitazone is profoundly affected. Noteworthy, this pathway is functional and was shown to respond to Pioglitazone under similar condition [49] in fibroblasts from human suffering from other pathological conditions such as mitochondrial infantile encephalomyopathy linked to Aifm1 mutation [50].

**Figure 4.**
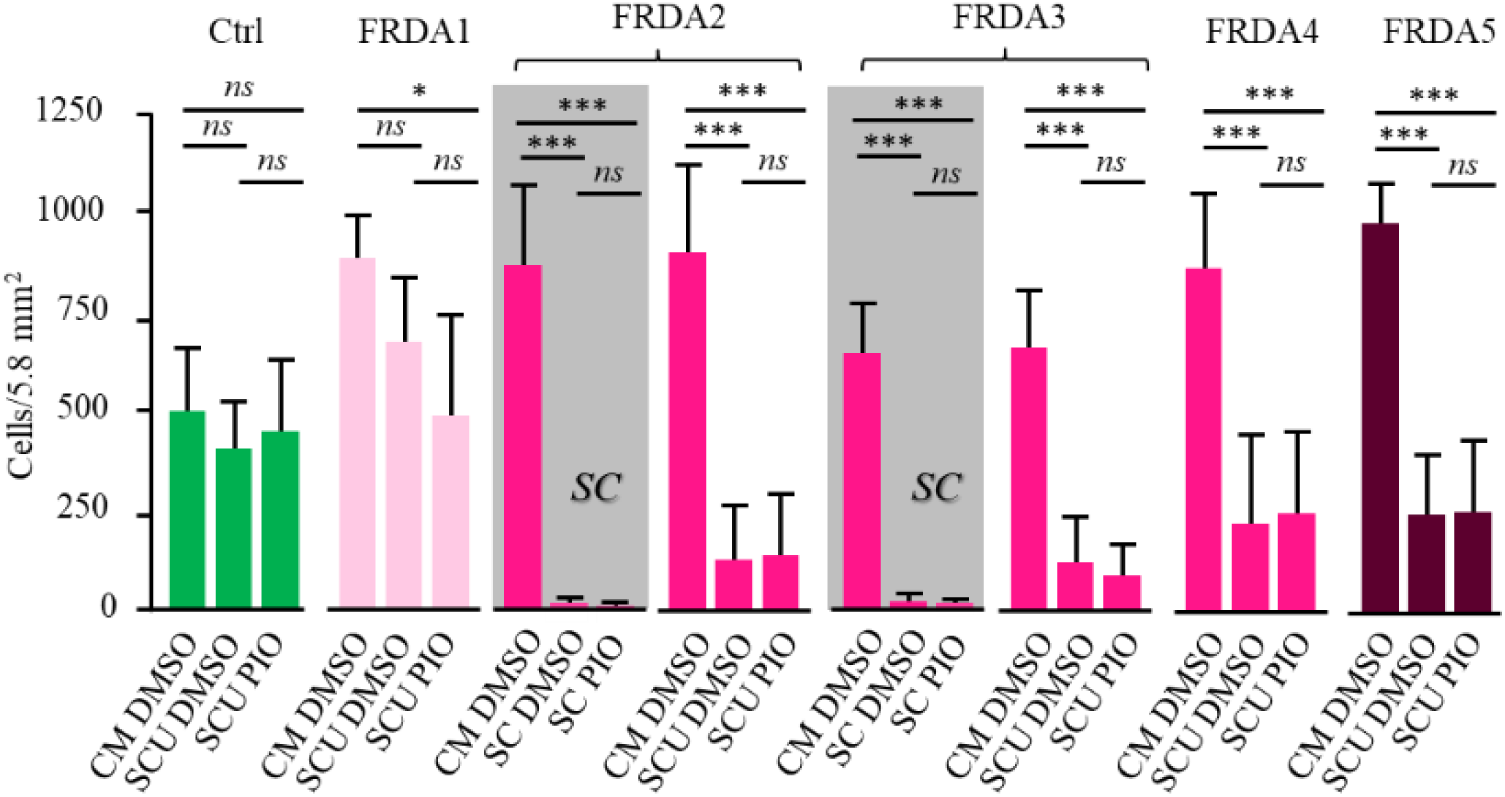
Proliferation of control (n=4; green) and FRDA fibroblast cells (FRDA1 to 5; red) in complete medium in the presence of DMSO (CM DMSO), under stressful conditions in the presence of DMSO and Uridine (SCU DMSO), or under stressful condition in the presence of Uridine and Pioglitazone (SCU PIO). In the two grey boxes, representative sets of data (obtained with FRDA2 and 3 fibroblasts) illustrate the total inefficiency of Pioglitazone to rescue growth of FRDA fibroblasts. Similar inefficiency was observed in the presence of Uridine for the five FRDA cells (FRDA1-5). Notice that added Pioglitazone was resuspended in DMSO. DMSO (0.001% final) was similar under the three conditions compared. Color code as described in Figure 2 legend. *ns*, non-significant.

### Correlation between sensitivity to oxidative stress and PIP_5_K_1_β mRNA expression

We next examined the potential correlation between mortality rate in FRDA fibroblast cultures grown in different media and the residual PIP_5_K_1_β mRNA in these cells (Table 2). Using cell survival as a criterion, 3 distinct types of behavior emerged. The first observed for patient 1 (FRDA 1) characterizes by a significant rescue by the addition of Uridine alone. Fibroblasts from this patient have been also the least impacted by stressing conditions. Interestingly enough, they have the highest Frataxin level and PIP_5_K_1_β mRNA expression. At the other end of the spectrum, fibroblast cultures from patient 5 were the most impacted by the stress conditions with less than 5% of the cell survival. Fibroblast culture from this patient was rescued neither by Uridine nor by Pyruvate alone. A partial survival could only be observed when both Uridine and Pyruvate were added (SCUP). Noticeably, patient 5 fibroblasts had the lowest PIP_5_K_1_β mRNA expression (Table 2). The growth of the three other patients could not be rescued by Uridine alone, while it was significantly improved by Pyruvate alone. Considering the recognized mechanism of Pyruvate acting through peroxide chelation, this suggests that impaired peroxide elimination, previously reported in fibroblasts from FRDA patients [11] was the likely cause of the poor survival of patient 2-5 fibroblasts under stressing conditions.

**Table 2.** Correlation between growth in the different culture media and PIP_5_K_1_β mRNA content. SC: stress condition (no Glucose, no Uridine, no Pyruvate); CM: complete medium with Glucose, Glutamine, Uridine, Pyruvate. SCU: SC *plus* Uridine; SCP: SC *plus* Pyruvate; SCUP: SC added with Uridine and Pyruvate. **+**: normal growth; **±**: intermediate growth; **-**: complete cell death at day 14. Concentrations used as indicated under Material and Methods.

| *Medium*<br>*Cells* | SC | SCU | SCP | SCUP | CM | PIP<sub>5</sub>K<sub>1</sub>β mRNA<br>(% of CTRL) |
| --- | --- | --- | --- | --- | --- | --- |
| FRDA1 | - | + | ± | + | + | 58 |
| FRDA4 | - | - | ± | + | + | 50 |
| FRDA5 | - | - | ± | ± | + | 43 |
| FRDA2 | - | - | ± | ± | + | 33 |
| FRDA3 | - | - | - | ± | + | 24 |

Quite noticeably, if the effect of Uridine, or Uridine plus Pyruvate on cell survival is judged by averaging the results obtained on cells from the five patients, it turns out to be statistically different (Fig.5). This was also observed in the case of fibroblasts from one FRDA patient (FRDA3) with a unexplained low catalase (Table. 1). Conversely, if repeated sets of data obtained for each patient are analyzed independently, it can be concluded that Uridine (FRDA1), or the simultaneous presence of Uridine and Pyruvate (FRDA1 to 4) exerts a statistically confirmed effect for 4 of the 5 patients (Fig.2;3).This clearly highlights the inter-individual variability and the impasse with statistics based on means.

**Figure 5.**
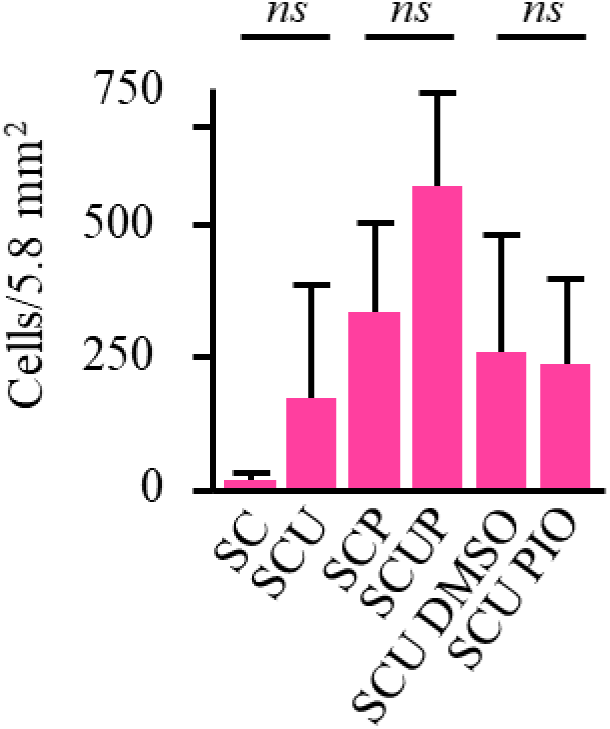
Averaging growth data for the five FRDA fibroblasts fully masks the protecting effect of Uridine (SC/SCU), or Uridine plus Pyruvate (SCP/SCUP), previously characterized for FRDA cells individually considered. When necessary (SCU DMSO/SCU PIO), DMSO (0.001% final) was similar. *ns*, non-significant.

## Discussion

This study reveals an unpredicted variability of cultured skin fibroblasts from patients harboring expansion in the frataxin gene. It is of course not envisaged here that cell cultures could reflect in any way the complexity of living organisms, especially human. As a result, their responses to the antioxidant agents cannot prefigure the results that might be subsequently observed on clinical trials. On the other hand, as is the case here, such studies can early point to inherent difficulties linked to the eminently variable consequences of even neighboring mutations, *i.e.* expansion in the Frataxin gene, when the effects of antioxidant agents are studied. They thus make it possible to discuss the difficulties encountered for more than 30 years in therapeutic trials using antioxidants to counter Friedreich’s Ataxia.

The study has been carried out under strictly controlled laboratory conditions and on a limited set of cells which would be usually described as homogeneous (all with large expansion, our inial selection criterion and PIP_5_K_1_β mRNA decrease). Yet, it reveals quite variable responses of cells to antioxidants. Most importantly, it first evokes the extreme variability of biochemical phenotypes, even when studying a single cell type (here, patient’s fibroblasts) from just five patients. Then it points to the place held by oxidative stress in this variability as seen through the response to three substances interfering with the oxidative stress. Indeed, this variability represents a hardly solvable difficulty in the management of clinical trials, as illustrated by the trial of Pioglitazone (Supplemental Fig. 2).

### The extreme variability of biochemical phenotypes

Most of research aiming at counteracting FRDA, including ours, attempt to identify efficient approaches or molecules to counterbalance the Frataxin mutation. For 30 years, several supposedly effective molecules, selected based on their indisputable effects in vitro or using so-called “models” of the disease (yeasts, *c. elegans*, *drosophila*, or mice) have been discarded because the real-life tests in human turned out to be disappointing, as none could reach sufficient statistical significance.

Several points must be considered when interpreting results from studies of models. First, as pointed out repeatedly in our study, variability is a fundamental characteristic of FRDA which undoubtedly stands, at least in part, from the genetic heterogeneity of patients. In contrast, most of the research models are constructed by introducing targeted genetic modifications which results in clonal populations thereby eliminating the essential genetic variability. These transgenic animals or cell lines are undoubtedly very valuable and irreplaceable tools for studying molecular mechanisms, but they are poor predictors of the efficacy or inefficacy of treatment. Indeed, to evaluate any therapeutic approach, one can hardly extrapolate the results obtained on clonal animal population as opposed to the genetically heterogeneous human population, or when bearing in mind the short lifespan of animal models while it takes several years, or even dozens for a disease to set in, or when taking into account the highly controlled and homogenous conditions in the animal facilities which cannot reflect a real life of patients exposed to variable and ever changing external factors. These factual concerns, which constitute real obstacles in carrying out therapeutic trials in humans, allows understanding the reason of the failures of numerous trials using substances deemed promising in the initial phases carried out on so-called models.

As mentioned above, a major limitation of these models results from the shortness of their life (at best 4-5 years) compared to the very progressive development of Friedreich’s disease. The reasons for this progressive character, observed in most mitochondrial diseases, are essentially unknown, but could turn out to be decisive in bringing about the eventual success or failure of any therapy. This point is far from being specific to Friedreich’s ataxia, having all its value for most mitochondrial diseases whose evolution is so poorly understood, and stands true for many complex human diseases.

### Oxidative stress can trigger death in FRDA skin fibroblasts cultures

Many biochemical abnormalities have been reported as the consequence of mutations in the Frataxin gene. They combine, depending on the tissue and possibly on individual, the deficit of iron-sulfur proteins [9], damage to the respiratory chain [9], impairment of oxidation of fatty acids [51], dysfunction of Pyruvate dehydrogenase [52], the abnormal levels of oxidative stress [10], the disorganization of cytoskeletal structures [12], glycolytic [53] or signaling pathways for antioxidant defenses [14]. By selecting fibroblasts for this study and selecting controlled stressing conditions resulting from strongly limited glucose availability and subsequent, limited source of reducing power (NADPH) for Glutathione known to control oxidative stress in FRDA cells [11,12], we can assess the cell lethality in response to oxidative stress and to compare the effect of compounds acting on oxidative stress and likely balancing these stressing conditions.

Although using one type of cells from only five patients, we observed a strong disparity between culture behavior and cells response to antioxidants. Stressful conditions had a devastating effect on all five patient’s cultures, but it could be significantly counterbalanced by Uridine only for patient 1 fibroblasts. In the presence of Pyruvate, the deleterious effect of stressing conditions was much less pronounced, being roughly annihilated and strongly reduced by Uridine for 4 of the 5 patients. Uridine and pyruvate have been described previously as neuroprotective compounds [33], our results confirmed their protective effects against oxidative stress in FRDA fibroblasts. Throughout this study, we were unable to evidence any positive effect of the glitazone (Pioglitazone) on the small number of FRDA fibroblasts tested.

Overall, it appears that the deleterious effect of oxidative stress and the protective effect of Uridine and/or Pyruvate on fibroblasts depend on each patient from which the culture originates.

### From culture flasks to clinical trials

Among the molecules tested here to illustrate the individual response to the oxidative stress of fibroblasts cultured under stressing conditions, Pioglitazone is worth discussing despite the lack of effect on cultured fibroblasts we have studied. Indeed, using this molecule to counteract Friedreich ataxia in human patients perfectly illustrate the difficulty in conducting human trials, when considering the variability between individuals, in the context of a rare disease such as FRDA.

Due to its ability to activate PPAR γ and cellular antioxidant defenses, this molecule was proposed for a proof of concept (Phase II) study in 2009. All the information on the procedures used in this trial is freely available (https://www.orpha.net/data/eth/GB/ID50165GB.pdf). The results of this trial have been made public, and it was stated that firm conclusion using the planned statistical analyzes is impossible to reach [54]. However, three lessons can be drawn from this trial now completed in France and which correspond to a recurrent problem in clinical trials in this disease. The first is related to the extreme heterogeneity among patients affected by this rare disease which makes it almost impossible to constitute sufficiently large cohorts allowing valid comparisons in studies carried out against placebo. A second finding relates to the duration of placebo effects. The careful management during the trials of patients who otherwise might feel to be left alone, without sufficient attention and hope for long period, results in some improvement in patients of the placebo group lasting even longer than a year. Importantly, the duration of many trials does not exceed such a long period of time. Finally, for the Pioglitazone trial, processing the data using a chosen Bayesian approach proved ineffective to obtain any convincing conclusion only allowing to state that the conditions were not met to seriously carry out this type of analysis. Indeed, this approach utilizes predictions made for the foreseeable outcome of the treatment for each patient which in the case of this disease, are based on unverified assumptions. Neither the profile of a patient at time t, nor his history prior to the start of the trial, sufficiently predicts the evolution of the disease with or without a treatment.

Summarized very briefly in Supplemental Fig. 2, the results indicate that on the basis of a composite score adding results from three scales for estimating the progression of the disease (ICARS [55], FARS [56] and SARA [57]) the score of a minority group of patients receiving Pioglitazone improved during the period of the test. In comparison, all patients receiving the placebo saw their composite score deteriorate or see inconsistent changes according to scale considered.

Thus, similarly to Idebenone, an antioxidant tested in rare mitochondrial diseases (FRDA and Leber’s hereditary optic neuropathy), the power of the trials was not, and probably will hardly be, sufficient to reach a confident statistical value [58]. In case of Idebenone, several clinical trials have shown a delay of neurological deterioration in FRDA disease in about 50% of the patients. Noticeably, the first publications focused on cardiac issues of a series of young patients report a reduction of cardiac hypertrophy and an improvement of cardiac functions resulting from the administration of Idebenone correlated with a recovery of enzymatic activities of iron-sulfur proteins, but did not at that time mention positive effect on neurological symptoms [59–64].

## Conclusion

To get out of this tricky situation, the way might be to stratify the patients better to perform trials and analyze their results. In the absence of reliable biomarkers, we need to find consensual criteria not reduced to a clinical picture at a given time. In that regard, distributing a few dozen people in placebo and treated groups has shown its limits in the case of FRDA over the past 30 years. The reality is that in the case of a rare, slowly progressive, and so heterogeneous disease like Friedreich’s ataxia, the group size will always be insufficient to allow standard tests to reach credible statistical values. As discussed above, even the Bayesian approach, as used for the Pioglitazone trial in France, has shown its limits. These latter could be due to our incapacity based on pre-trial progression of the disease to produce credible predictions for each individual disease evolution.

Defining the criteria is a risky business. These criteria should include a maximum of parameters likely to determine the expression of the disease. Beside detailed clinical parameters, molecular and biochemical parameters might/should include size of the Frataxin expansion, respiratory chain and Krebs cycle activities, oxidative stress intensity, extent of PIP_5_K_1_β silencing, etc. These are all related to the primary FRDA mutation and supposedly determining for clinical expression of the disease. As illustrated for skin fibroblasts in our study, the impairment of the expression of the PIP_5_K_1_β expression might be one of these criteria and failure to consider this parameter can be sufficient to mask the positive effect of a cell treatment. As for last decades, the distribution of quite limited number of patients affected by this rare disease between treated- and placebo-groups will likely lead to inconclusive data or in conclusions biased by a given sampling. In particular, this greatly increases the risk of being unable to conclude on the usefulness of any molecule or approach trialing and to throw the baby (promising molecules or strategies) out with the bath water.

Here, the study of FRDA fibroblasts was useful to stress the major inter individual differences between patient’s cells toward oxidative stress and their responses to antioxidants, presumably associated with different genetic background. In the future, the use of the powerful tools now available for omics analyses (genomics, transcriptomics, proteomics, metabolomics) might disclose one or more rational that could account for the observed differences between FRDA individuals at cellular and organismal levels. At cellular level, this rather expensive investigations should be better undertaken using more appropriated cell types (neurons, cardiomyocytes) known to be involved in the disease. This panel of omics which should be completed by past and present “environmental omics” when dealing with patients. So far however omics approaches did not provided markers strong enough to be used in clinical trials.

## Funding

General expenses were covered by grants from AAJI (Association pour l’Aide aux Jeunes Infirmes & aux Personnes Handicapées), AFAF (Association Française de l’Ataxie de Friedreich) and Association OLY (Ouvrir Les Yeux).

## Author Contributions

Conceptualization: Paule Bénit. Funding acquisition: Paule Bénit, Pierre Rustin. Investigation: Paule Bénit. Methodology: Paule Bénit. Supervision: Pierre Rustin, Malgorzata Rak. Writing – original draft: Paule Bénit. Writing – review & editing: Pierre Rustin, Malgorzata Rak.

## Competing interests

The authors have declared that no competing interests exist.

## Abbreviations

FRDA: Friedreich Ataxia
CM: Complete Medium
SC: Stressing Condition
SCU: Stressing Condition *plus* Uridine
SCP: Stressing Condition *plus* Pyruvate
SCUP: Stressing Condition *plus* Uridine and Pyruvate

## Notes

### Competing Interest Statement

The authors have declared no competing interest.

