## supplemental fig 1 and 2 for "One-to-one Benefit provided by Antioxidants to cultured skin Fibroblasts from Friedreich Ataxia patients"

**Supplemental material**

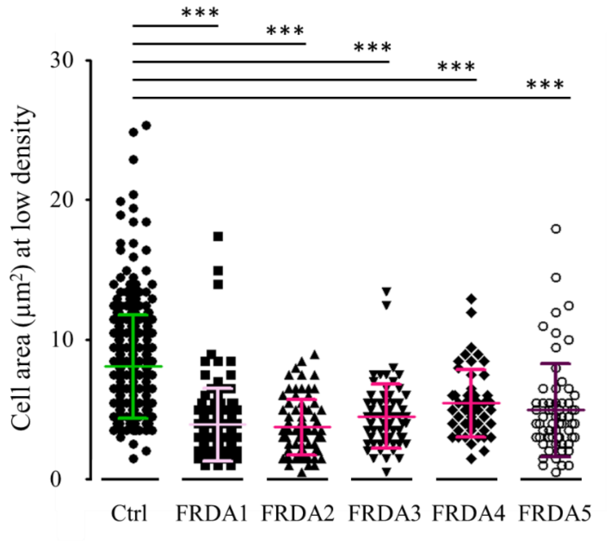


Supplemental Figure 1. Comparison of control and FRDA fibroblast cell areas. Cell area was estimated from random image sampling and found much reduced for FRDA fibroblasts.

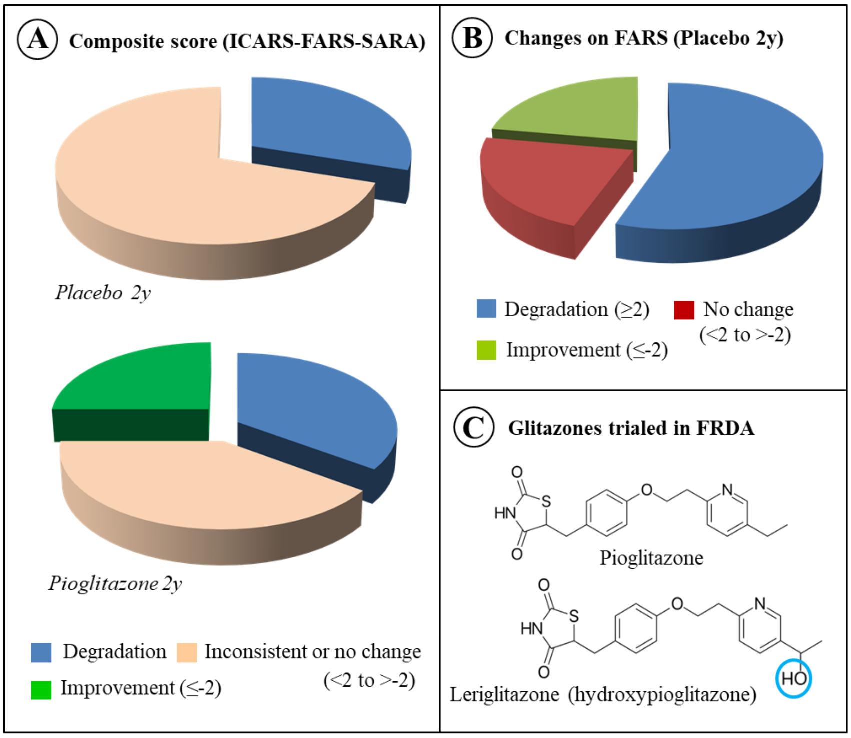


Supplemental Figure 2. **State of art for the use of glitazone to counteract Friedreich ataxia**. **A**, Effect of Pioglitazone *versus* placebo on the composite score resulting from the integration of results from three scales: International Cooperative Ataxia Rating Scale (ICARS), Functional Assessment Rating Scale (FARS), Scale for Assessment and Rating Ataxia (SARA). Changes ≥ 2 points were taken as significant. This clinical trial was carried out at Robert Debré Hospital (Paris, France) on FRDA patients starting 2008. Details of the trial conditions are available at https://www.orpha.net/data/eth/GB/ID50165GB.pdf, while conclusions tentatively drawn from the trial have been previously published [54] **B,** The spectacular duration (2 years) of the placebo effect, traced by the significant number of patients displaying an improvement when considering the FARS scale alone (Friedreich Ataxia Rating Scale). **C**, another member of the glitazone family, Leriglitazone (hydroxyPioglitazone), has been reported to be beneficial against FRDA in a phase 2 proof of concept (and a phase 2/3 clinical trial in another central nervous system disorder, adrenomyeloneuropathy) in US (https://www.minoryx.com/media/minoryx’s_clinical_candidate_leriglitazone_shows_clinical_benefit_in_a_proof_of_concept_phase_2_study_in_friedreichs_ataxia/).
